# Behaviorally prioritized entity-relation structure captures human visual cortical representations of natural scenes

**DOI:** 10.64898/2026.09.16.752038

**Authors:** Yichen Wu, Wenyi Jiang, Sheng Li

## Abstract

Understanding natural scenes requires identifying visible entities and representing how those entities are related. Recent studies have shown that artificial neural networks (ANNs), large language models (LLMs), and vision language models (VLMs) can predict visual cortical responses to natural images. However, the neural organization of relational scene meaning remains poorly understood, in part because these models typically encode scene content in global feature spaces that are difficult to decompose into separable entity and relation components. Here, we combined scene-graph annotations, behavioral measurements, and large-scale neural datasets to characterize structured relational representations during natural vision. We used RotatE, a knowledge-graph embedding model, to represent head-relation-tail triplets annotated for images from the 7T Natural Scenes Dataset. Triplet embeddings reliably captured cortical representational structure across the visual hierarchy. Behavioral judgments further revealed systematic differences in triplet accessibility associated with visual, relational, and graph properties. Prioritizing more behaviorally accessible triplets improved neural correspondence and explained unique variance beyond object co-occurrence, ANN image features, and LLM caption embeddings. Decomposing triplet representations into entity and relation components revealed partially dissociable cortical contributions, with lateral parietal cortex showing sensitivity to both. Triplet-based semantic information also remained spatially grounded: visual-field-specific triplet models preferentially predicted voxels with matching retinotopic preferences. Finally, cross-species comparison indicated that triplet-based semantic features were relatively more aligned with human high-level visual cortex than with macaque inferotemporal cortex. Together, these findings provide new insights into the representation of semantic relational information in the human visual cortex during natural scene perception.

## Introduction

Natural vision conveys more than a list of visible entities. To understand a scene, observers must also recover how objects, people, animals, and background elements are related. These relations specify spatial configurations, actions, affordances, and social interactions that cannot be inferred from entity identity alone. A representation that encodes only which entities are present would therefore omit a substantial part of what makes a natural scene meaningful.

Despite the central role of relational information in scene understanding, its representational organization in the human brain remains poorly characterized. Traditional neuroimaging studies have extensively examined category-and object-selective responses in high-level visual cortex (Grill-Spector & Weiner, 2014; Peelen & Downing, 2017; Ritchie et al., 2026). More recently, artificial neural networks (ANNs), large language models (LLMs), and vision–language models (VLMs) have substantially improved the prediction of neural responses to natural images (Bogdan et al., 2026; Doerig et al., 2025; Guclu & Van Gerven, 2015; Rong et al., 2025). In particular, caption-based language models use sentences that summarize scene meaning as input, thereby incorporating both entity and relation information. However, these models represent an entire caption as a single global semantic embedding vector, making it difficult to determine the specific contributions of entities and relations. Pixel-based ANNs may also implicitly encode semantic information in their high-layer representations, but their representations face the same limitation.

This limitation is important because relations may constitute a representational component that is at least partly separable from the entities they connect (Holyoak et al., 2022). Recent studies using controlled semantic judgments have shown that relation information can generalize across different concept pairs and can be distinguished from concept information in distributed cortical regions (Chen et al., 2026). Yet it remains unclear how such relational structure is represented during natural vision, where multiple entities and multiple relations coexist within the same image. Global pixel-based or caption-based embeddings generally entangle these components, limiting their ability to reveal how relation information is internally organized in scene representations.

A second challenge is that the semantic information available in a natural image is not necessarily equivalent to the information that observers actually process. A complex scene may contain many entities and relations, but semantic information is not equally available across the image (Haskins et al., 2026; Henderson & Hayes, 2017). Entities and relations at different spatial positions, or with different meanings, may differ in their perceptual accessibility (Henderson & Hayes, 2017; Kaiser & Cichy, 2018; Murlidaran & Eckstein, 2025; Zhao & Koch, 2013). The pixel-and caption-based models may therefore incorporate all available semantic information, even though only part of that information may strongly contribute to the neural response evoked during image viewing. Using behavioral measures to determine which entities and relations are perceptually accessible would help identify the subset of semantic information most likely to shape neural representations during natural viewing.

Scene graphs offer a structured representation that is well suited to addressing these questions (Chang et al., 2023; H. Li et al., 2024). A scene graph describes an image as a set of relational triplets of the form *head-relation-tail*, such as *person-riding-horse* or *cup-on-table*. As the basic units of a scene graph, triplets provide a more structured description of scene meaning than object labels while remaining more explicit than global captions. Each triplet specifies two entities and the relation between them, thereby preserving relational information without collapsing the image into a single semantic summary. Moreover, because the entities can be localized to specific regions of the image, the triplet also carries local spatial information. This makes triplets useful for investigating which components of a scene’s semantic structure are perceptually accessible and where those components are represented in cortex. Instead of asking subjects to freely describe an image or interpret a whole caption, we can ask whether a specific triplet was perceived in a specific image.

Here, we hypothesized that human visual cortex represents natural scenes through a behaviorally prioritized set of relational triplets. We used RotatE, a knowledge-graph embedding model, to derive quantitative representations of scene entities and relations (Sun et al., 2019). We first asked whether these triplet-based representations predict neural activity elicited by natural scenes. We then used a triplet judgment task to determine which semantic information was consistently perceived across subjects, and whether weighting the model by perceptual accessibility improved its predictive power. Leveraging the decomposable and spatially localized nature of the triplet representation, we further examined the distinct contributions of entities and relations and tested whether representations of local relational content conform to the retinotopic organization of high-level visual cortex. Finally, using neural responses to the same natural images in macaque inferotemporal cortex, we asked whether the relative contribution of triplet-based semantic structure differs between human and nonhuman primate visual systems. Together, these analyses provide a structured account of how relational scene information is selected and organized in the human brain.

## Results

We first investigated how relational triplets are represented in the human visual system using the Natural Scenes Dataset (NSD), a large-scale public 7T fMRI dataset (Allen et al., 2022). Subjects in NSD viewed a large set of natural scene images, for which triplet annotations were obtained from the Panoptic Scene Graph (PSG) dataset (Yang et al., 2022). We used RotatE to quantitatively transform the triplets into vector-based embeddings. In RotatE, each relation is defined as a rotation from head vector to tail vector in a complex-valued vector space (Figure 1A). We trained the RotatE model on the triplet annotations from PSG dataset, yielding a 600-dimensional embedding vector for each *head-relation-tail* triplet, with 200 dimensions for each component (Figure 1B). These embeddings provided a quantitative link between the semantic triplets of the images and the neural responses evoked when subjects viewed those same images.

**Figure 1.**
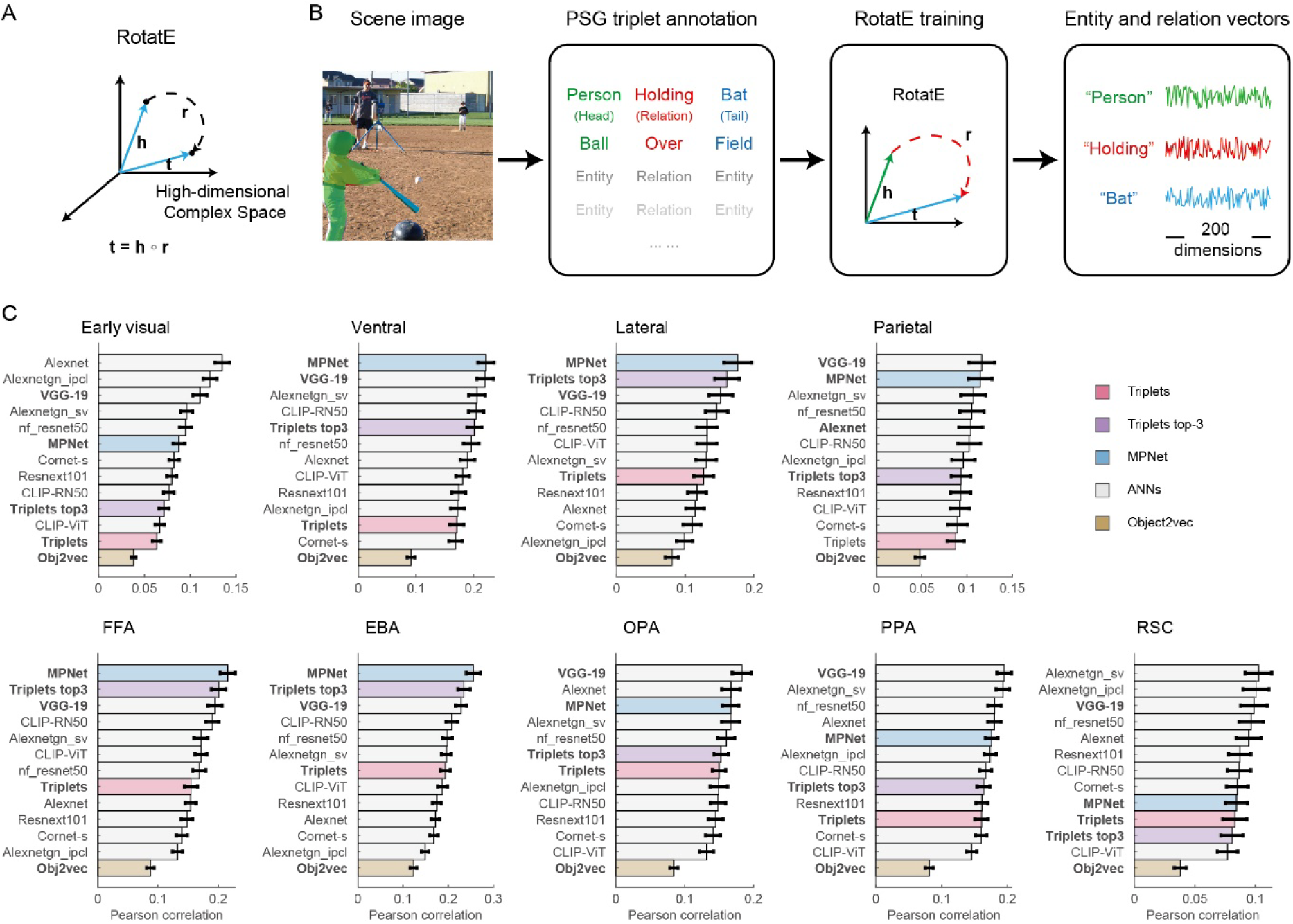
RotatE model and its prediction of visual cortical activity. (A) Schematic of the RotatE model. The head (h) and tail (t) entities of a triplet are projected as vectors in a high-dimensional complex-valued space, whereas the relation (r) defines an element-wise rotation. During training, the model learns embeddings such that the tail vector approximates the head vector after rotation by the relation. (B) From natural scene images to RotatE vectors. In the PSG dataset, each image is annotated with a set of triplets. These triplets are used to train RotatE and derive vectors for the corresponding entities and relations. Each 200-dimensional vector shown here was obtained by concatenating the real and imaginary components of its complex-valued vector. (C) ROI-wise comparison of RSA model performance. The top row shows results for the four visual stream ROIs, and the bottom row shows results for the five category-and scene-selective ROIs. Models based on different input types and architectures are ordered according to their ability to predict neural activity in each ROI. The models shown in bold were used in the statistical comparisons reported in the main text.

### Triplet embeddings predict visual cortical representations

We averaged the embeddings of all triplets within each image to derive an image-level triplet embedding. These image-level embeddings were used to construct a triplet-based representational dissimilarity matrix (RDM), which was compared with each subject’s fMRI response RDM using representational similarity analysis (RSA). Triplet-model RSA correlations were significantly greater than zero in all ROIs (all *q* <.001, FDR corrected; Figure 1C), including early visual cortex; the ventral, parietal, and lateral visual streams (using the NSD’streams’ ROI definitions); and higher-level category-and scene-selective regions, including the FFA, EBA, OPA, PPA, and RSC.

We next compared the triplet model with models that characterize image semantics using different inputs and architectures. These included an object2vec model (Bonner & Epstein, 2021) based on co-occurrence statistics of NSD panoptic labels, an ANN model (VGG19) trained to classify image category from pixel values, and an MPNet model (Doerig et al., 2025) that use scene captions as input. Planned contrasts from repeated-measures ANOVAs, with ROI and model as within-subject factors, showed that, across the three visual stream ROIs, the triplet model outperformed object2vec (Δ = 0.055, *q* <.001) but performed less well than VGG19 (Δ =-0.034, *q* <.001) and MPNet (Δ =-0.043, q <.001). The same pattern was observed across FFA, EBA, PPA, OPA, and RSC: triplets exceeded object2vec (Δ = 0.066, *q* <.001), but were lower than VGG19 (Δ =-0.032, *q* <.001) and MPNet (Δ =-0.031, *q* <.001).

Together, these results indicate that the triplet model provides an effective and interpretable account neural representations across high-level visual cortex. Its advantage over object2vec highlights that incorporating relational structure captures information not available from entity co-occurrence alone. Moreover, the decomposable and spatially localized structure of triplets makes it possible to relate specific entities and relations to their corresponding locations within an image. We therefore hypothesized that the interpretability of the model would be increased by prioritizing triplets that were more likely to be perceptually accessible to human observers during image viewing.

### Features determining the perceptual accessibility of triplet

We conducted a behavioral experiment to identify the factors that determine which triplets are most likely to be perceived. All subjects performed a triplet judgment task involving the same set of 720 triplets drawn from 240 images. For each image, subjects were asked to identify the incorrect components (i.e., head, relation, or tail) among three triplets that were inconsistent with the image (Figure 2A). Inter-subject reliability was quantified as the mean pairwise proportion of identical responses to the same triplets, yielding a reliability of 0.6655. A permutation test with 1,000 iterations showed that this agreement was significantly above chance (*p* <.001), indicating reliable shared tendencies across subjects in the triplets and triplet components they perceived (Figure 2B).

**Figure 2.**
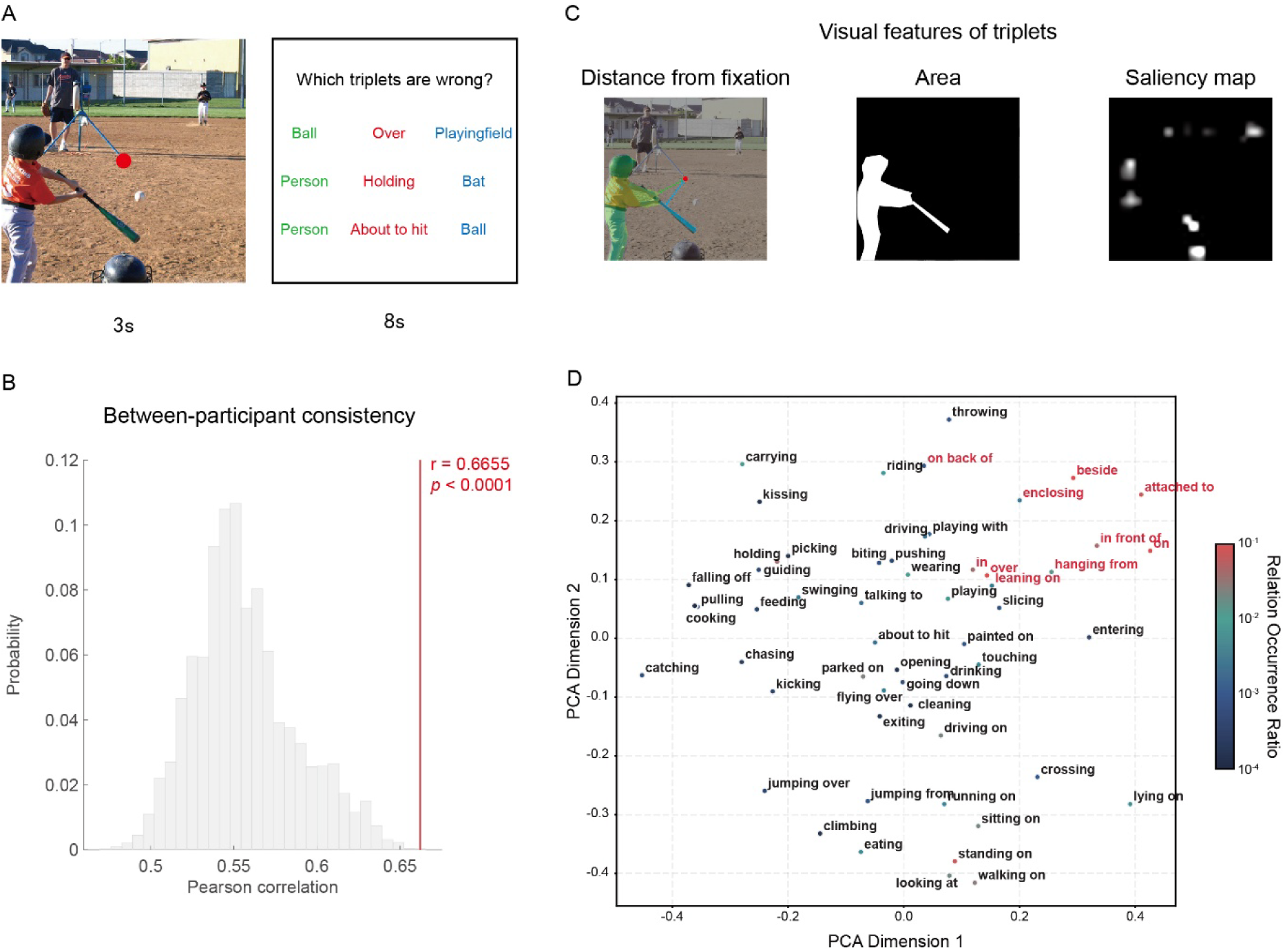
Behavioral assessment of the perceptual accessibility of scene triplets. (A) Experimental procedure. In each trial, subjects first viewed a scene image while maintaining fixation on a central red dot. They were then presented with three triplets and asked to judge which components of the three triplets were incorrect. (B) Inter-subject consistency in triplet judgments. The gray histogram shows the permutation distribution of the mean pairwise proportion of identical responses across subjects, generated from 1,000 iterations. The red line indicates the observed inter-subject consistency. (C) Visual features used to characterize the perceptual accessibility of triplets. (D) PCA visualization of relation vectors. Relation vectors are projected onto the first two principal components. Red labels denote geometric relations, and black labels denote non-geometric relations. The color of each point indicates the frequency of the corresponding relation in the full PSG dataset.

To determine why some triplets were more likely to be perceived than others, we defined a set of triplet features and examined their ability to predict the behavioral performance in the triplet judgment task. First, we expected the visual features of a triplet to influence its perceptual accessibility. We therefore included features describing the triplet’s retinal location and the visual prominence of its entities, including distance from central fixation, entity area, and visual saliency (Walther & Koch, 2006) (Figure 2C).

Second, we expected the semantic content and predictability of a triplet to influence its perceptual accessibility. Principal component analysis (PCA) of the relation embeddings showed that geometric relations clustered in the principal component space, suggesting that the distinction between geometric and non-geometric relations constituted a meaningful feature of the embeddings (Figure 2D). Geometric relations, such as *on*, *in front of*, and *attached to*, primarily describe the relative spatial configuration between two entities, whereas non-geometric relations convey richer semantic or action-based information, as exemplified by *crossing* or *carrying*. In addition to relation type, we included triplet score and semantic alignment as relational features. Triplet score indexes the plausibility of a triplet and was defined as the negative distance between the rotated head embedding and the tail embedding. Semantic alignment was defined as the cosine distance between the triplet embedding and the whole-image embedding, indexing how well the triplet fits the broader image context.

Third, we included graph-related features characterizing the position of each triplet within the scene graph, including triplet count and triplet degree. These features captured the frequency with which a triplet appeared in the scene and the extent to which its constituent entities were connected to other nodes.

A logistic mixed-effects model showed that visual, relational, and graph-based features predicted subjects’ behavioral responses. Specifically, false alarms were higher for geometric than for non-geometric relations (β =-0.348, *t* =-6.03, *p* <.001), higher for triplets located farther from fixation (β = 0.124, *t* = 5.60, *p* <.001), lower for triplets with larger entity areas (β =-0.292, *t* =-11.25, *p* <.001), lower for more visually salient triplets (β =-0.070, *t* =-3.40, *p* =.001), and lower for triplets that appeared more frequently in the scene graph (β =-0.128, *t* =-4.49, *p* <.001). The full results of the logistic mixed-effects model are reported in Supplementary Table 1.

Together, these results show that triplets differed systematically in their perceptual accessibility and that this variability was related to their visual properties, relation type, and frequency of appearance. These findings provide a behavioral basis for refining the triplet model: rather than treating all triplets within an image as equally informative, the neural model can assign greater weight to triplets that are more likely to be perceived by human observers.

### Behaviorally prioritized triplet model improves neural prediction

We next constructed behaviorally prioritized triplet model for RSA (Figure 3A). Specifically, we refitted a reduced logistic mixed-effects model that included the three visual features (distance from fixation, entity area, and visual saliency) as well as relation type (geometric and non-geometric) (Figure3A and Supplementary Table 2). The fitted coefficients were then used to compute a behavioral weight for each triplet (Figure 3A, middle). We ranked the triplets within each image according to these weights, constructed image-level embeddings by averaging the top 1 to 6 triplets, and evaluated the neural predictivity of each model with RSA. As shown in Figure 3B, the optimal number of behaviorally selected triplets varied across visual regions. In the ventral stream, lateral stream, FFA, and EBA, embeddings based on the top 3 triplets yielded the highest model performance. In the parietal stream and the three scene-selective regions, including additional triplets produced further but relatively modest improvements. We therefore selected the behaviorally prioritized top-3 triplet model for subsequent analyses.

**Figure 3.**
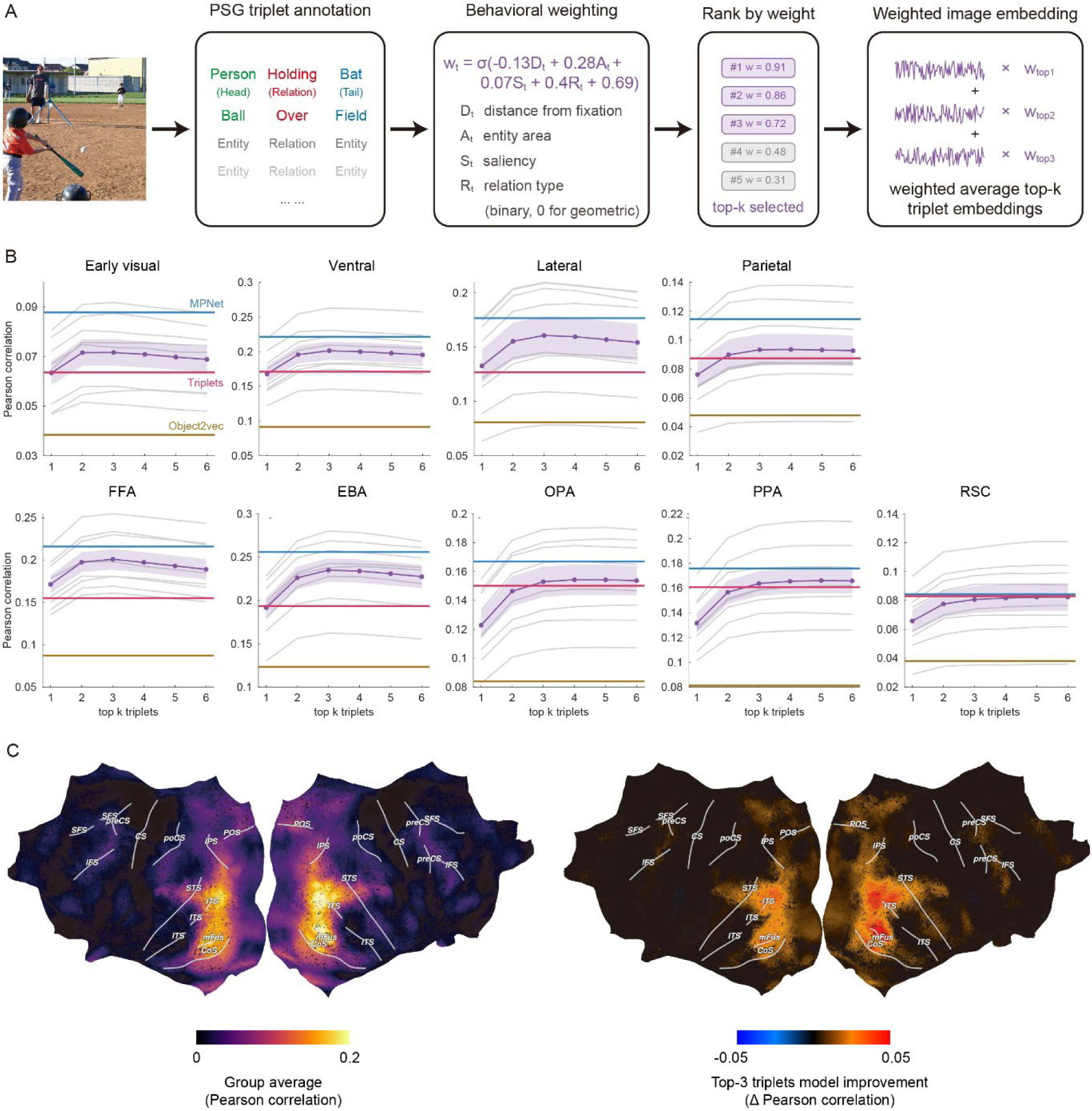
Behaviorally prioritized triplet representations improve neural prediction. (A) Construction of the behaviorally prioritized top-k triplet model. Triplet annotated in PSG dataset were first assigned behavioral weights based on their visual and relational features. Within each image, triplets were ranked according to these weights, and the embeddings of the top-k triplets were averaged to obtain an image-level embedding. (B) Effects of the number of selected triplets on ROI-wise neural predictivity. The blue, red, and yellow horizontal lines indicate the mean RSA performance across eight subjects for MPNet, the unweighted triplet model, and obj2vec, respectively. The purple line and shaded region indicate the mean and standard error of the top-k triplet model, respectively. The light gray lines indicate the performance of individual subjects. (C) Searchlight RSA results. The left map shows the group-averaged Pearson correlation between the top-3 triplet RDM and neural RDMs. The right map shows the difference in RSA correlations between the top-3 triplet model and the unweighted triplet model, highlighting cortical regions in which behavioral prioritization improved neural prediction. Statistical significance was assessed using right-tailed t tests against zero with Storey’s FDR correction. Non-significant vertices are shown in black. Abbreviations: CoS, collateral sulcus; CS, central sulcus; IFS, inferior frontal sulcus; IPS, intraparietal sulcus; ITS, inferior temporal sulcus; mFus, mid-fusiform sulcus; poCS, postcentral sulcus; POS, parieto-occipital sulcus; preCS, precentral sulcus; SFS, superior frontal sulcus; STS, superior temporal sulcus.

Across the three visual stream ROIs, the top-3 triplet model outperformed the unweighted triplet model (Δ = 0.023, *q* <.001), although its performance remained lower than that of VGG19 (Δ =-0.011, *q* = 0.002) and MPNet (Δ =-0.013, *q* <.001). The same pattern was observed across FFA, EBA, PPA, OPA, and RSC: the top-3 triplet model outperformed the unweighted model (Δ = 0.018, *q* <.001) but performed less well than VGG19 (Δ =-0.014, *q* <.001) and MPNet (Δ =-0.013, *q* <.001). These results demonstrate that prioritizing perceptually accessible triplets improves neural prediction, suggesting that semantic processing of scene triplets significantly influence the visual cortical responses.

We next asked whether the triplet model explained neural representations beyond information captured by other existing models. We conducted partial correlation analyses including the top-3 triplet model, object2vec, VGG19, and MPNet. For each model, we tested whether it accounted for unique variance in the neural RDMs after controlling for the other three models (Supplementary Figure 1). Partial correlations for the top-3 triplet model were significantly greater than 0 in all nine ROIs (all *q* <.001, FDR corrected), indicating that behaviorally prioritized triplets capture aspects of neural representational structure that is not reducible to object co-occurrence, scene captions, or high-level CNN features.

Having established the unique contribution of behaviorally prioritized triplet representations, we next examined where these representations were expressed across the cortical surface. Searchlight RSA allowed us to characterize the cortical distribution of triplet-based semantic information, both within the visual hierarchy and the regions involved in high-level conceptual and relational processing.

The top-3 triplet model predicted neural activity across the visual hierarchy, with substantially stronger effects in higher-level visual regions than in early visual cortex (Figure 3C, left). Significant effects were also observed beyond visual cortex, including in lateral parietal cortex (LPC), medial parietal cortex (MPC), and prefrontal cortex (PFC), regions commonly implicated in conceptual and relational processing. A direct comparison with the unweighted triplet model further showed that behavioral prioritization improved neural prediction across high-level visual cortex, with particularly pronounced gains near LPC and fusiform sulcus (Figure 3C, right).

### Cortical representations of entities and relations

Scene triplets offer a structured means of dissociating representations of entities from representation of the relation between them. To examine these components separately, we decomposed each image-level triplet embedding into a relation component and a combined head-tail entity component. We constructed separate RDMs for the relation and entity components, and calculated partial Pearson correlations to estimate the unique contribution of each component while controlling for the other. Searchlight analysis revealed unique entity-related information in inferior temporal cortex and areas along the intraparietal sulcus (IPS), whereas LPC showed evidence for both entity and relation representations (Figure 4A). ROI-level partial-correlation analyses revealed a similar pattern, further supporting a distinction between the cortical distributions of entity and relation information (Figure 4B).

**Figure 4.**
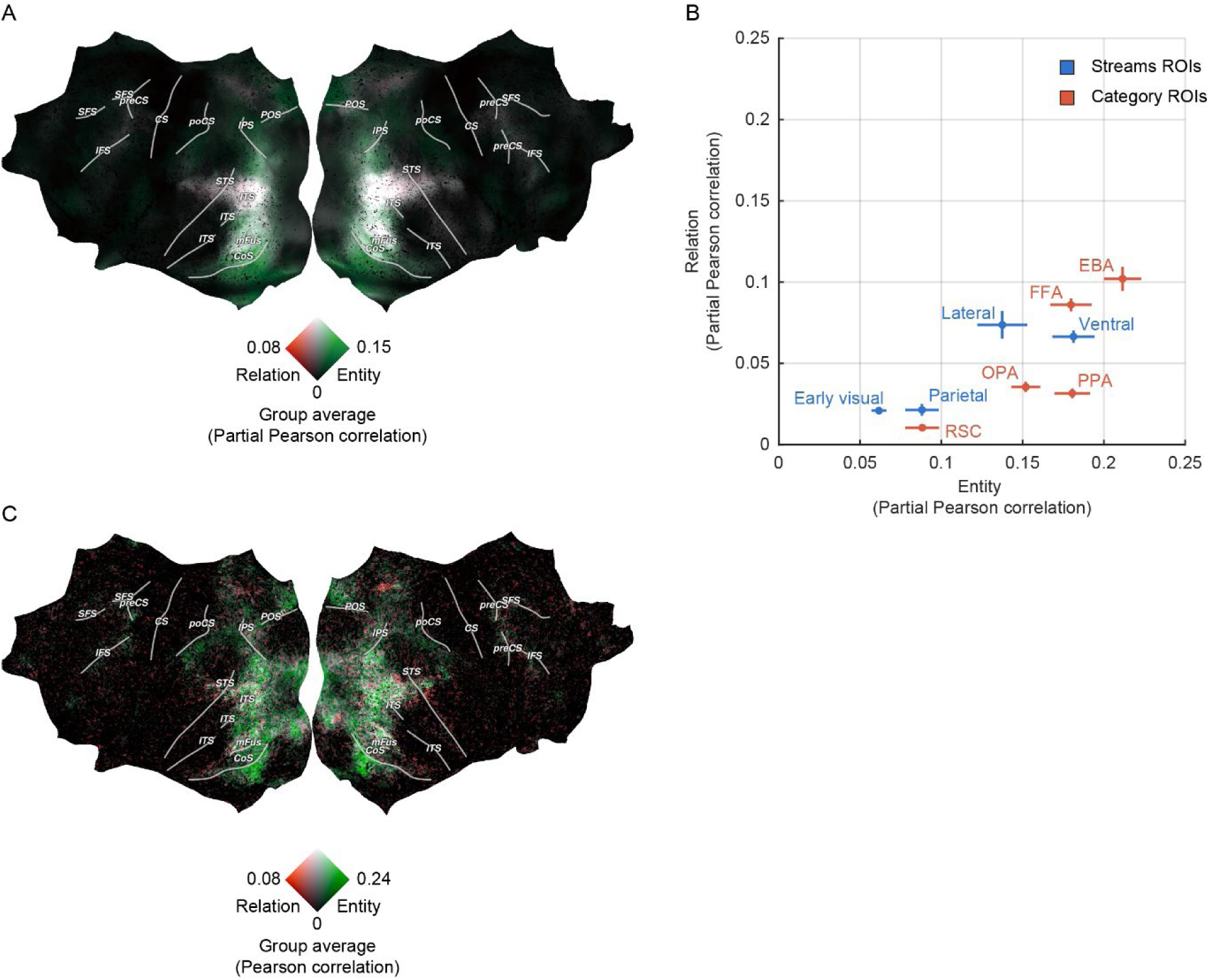
Cortical representations of entities and relations. (A) Searchlight RSA of entity and relation representations. The maps show partial Pearson correlations for entity and relation embeddings, computed within vertices that were significant in the searchlight analysis shown in Figure 3C. (B) ROI-level representations of entities and relations. Partial Pearson correlations quantify the unique contribution of entity and relation embeddings. Points represent subjects’ mean partial correlations within each ROI, and error bars indicate the standard deviations across subjects. (C) Encoding model results. The maps show respective contributions of entity and relation features to encoding model performance, computed within vertices showing significant prediction in the encoding analysis shown in Supplementary Figure 2.

We further used voxel-wise encoding models to predict neural activity from the entity and relation components of the triplet embeddings. Consistent with the searchlight RSA results, both components contributed to prediction of neural activity in LPC (Figure 4C, see Supplementary Figure 2 for encoding model results based on image-level triplet embeddings). The converging results indicate that entity and relation information make partially dissociable contributions to cortical representations of natural scenes, while converging in the LPC.

### Triplet representations are spatially grounded and context-independent

The partially dissociable neural representations of entities and relations demonstrate one source of the triplet model’s interpretability. Two additional properties of its representational structure, spatial localization and context-independence, further enable the model to generate specific predictions about cortical scene representations.

Triplet embeddings are local: each selected triplet can be spatially localized to the image regions occupied by its constituent entities. This spatial structure allowed us to test whether triplet-based semantic representations were aligned with retinotopic preferences in high-level visual cortex. For each triplet, we multiplied its behavioral weight by the proportion of its entity regions falling within the left or right visual field. Triplets were then reranked separately for each visual field to construct left-and right-visual-field top-3 triplet models. Using population receptive field (pRF) estimates, we divided the voxels within each ROI into groups preferring the left or right visual field. Across ROIs, voxels preferring a given visual field were better predicted by the top-3 triplet model constructed from the matching visual field than by the model constructed from the opposite visual field (Figure 5A). Reliable model x visual-field interactions were revealed in separate ROI x model x visual-field ANOVAs for the visual stream ROIs (*F* = 37.30, *p* <.001) and the high-level category-and scene-selective ROIs (*F* = 68.78, *p* <.001) (Figure 5C).

**Figure 5.**
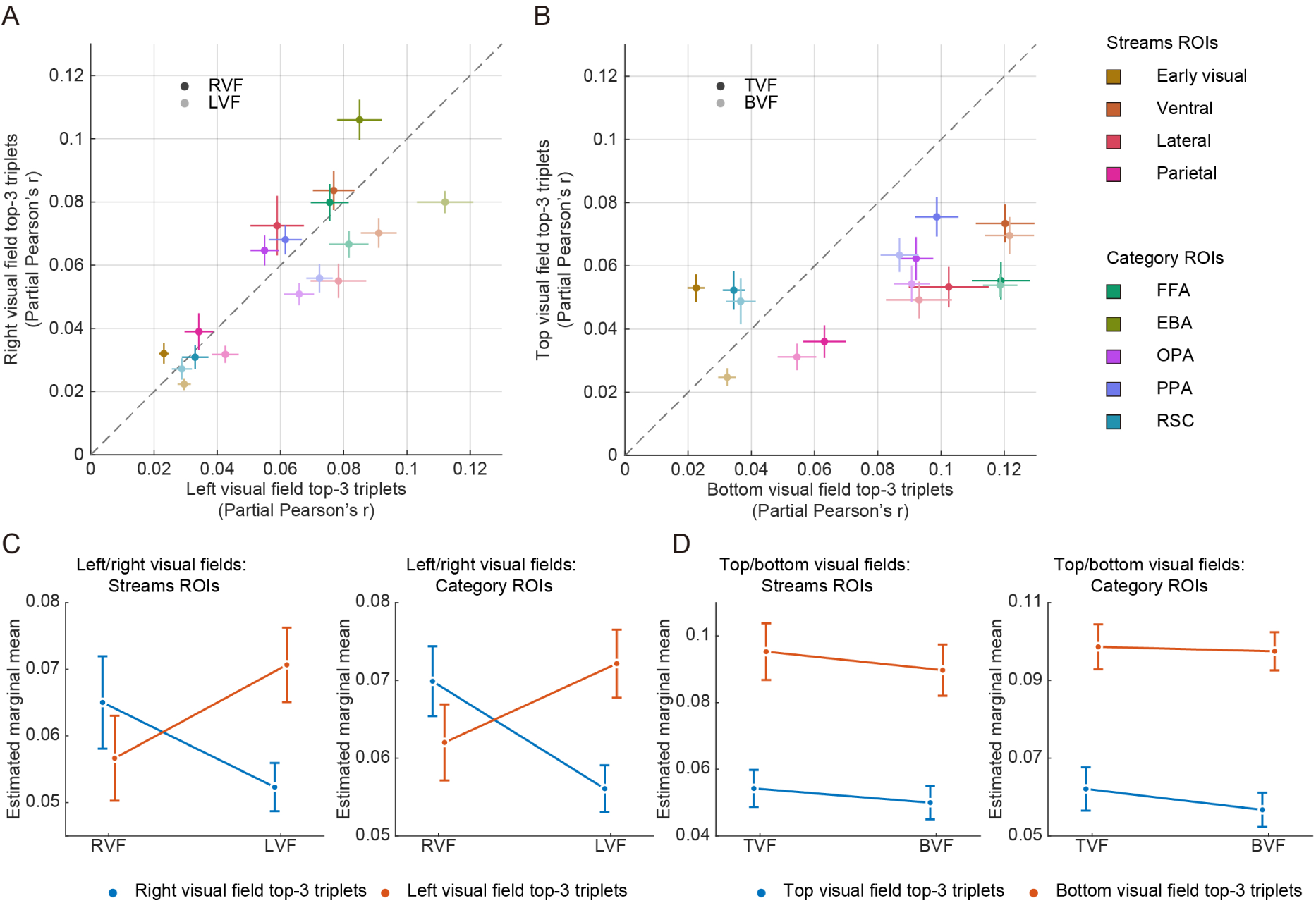
Visual-field-specific neural predictions from top-3 triplet model. (A) Visual-field-specific neural prediction along the horizontal dimension. Partial Pearson correlations quantify the correspondence of left-and right-visual-field top-3 triplet models with neural activity from voxels preferring the left (LVF) or right (RVF) visual field within each ROI. (B) Visual-field-specific neural prediction along the vertical dimension. Partial Pearson correlations quantify the correspondence of top-and bottom-visual-field top-3 triplet models with neural activity from voxels preferring the top (TVF) or bottom (BVF) visual field within each ROI. In (A) and (B), points indicate mean partial correlations across subjects within each ROI; horizontal and vertical error bars indicate the corresponding standard errors of the mean. (C) Marginal effects from the left/right visual-field ANOVAs, showing the model x visual-field interactions in visual stream ROIs and category-and scene-selective ROIs. (D) Marginal effects from the top/bottom visual-field ANOVAs, showing the main effects of triplet model in the two ROI groups.

Applying the same procedure to the top and bottom visual fields revealed a main effect of triplet model in both the visual streams ROIs (*F* = 116.9, *p* <.001) and the category-and scene-selective ROIs (*F* = 458.3, *p* <.001). Triplet models derived from the bottom visual field predicted neural activity more strongly than the top visual field (Figure 5B and 5D), consistent with the idea that the bottom visual field generally contained richer semantic information in natural scenes (Greene, 2013). Together, these results show that the relational summary captured by the top-3 triplet model is not only compact and behaviorally prioritized, but also spatially grounded in retinotopically organized cortical responses. More broadly, the spatially localize structure of triplet representations offers a means of investigating how spatial attention, fixation, and eye movements dynamically select relational information from natural scenes.

Although each triplet can be localized within an individual image, its entity and relation labels are encoded in a shared embedding space learned across triplets in all images. This context-independent structure allowed us to construct targeted exploratory contrasts probing the cortical organization of different types of scene semantics. After removing the entity components, we constructed separate embeddings for geometric and non-geometric relations and passed each through the trained encoding model to predict neural responses. We then contrasted the predicted responses for non-geometric versus geometric relations (Figure 6A). Stronger preference for non-geometric than geometric relations was observed near the LPC and fusiform cortex. These regions overlapped with those in which behavioral prioritization produced the largest improvement over the unweighted triplet model (Figure 3C).

**Figure 6.**
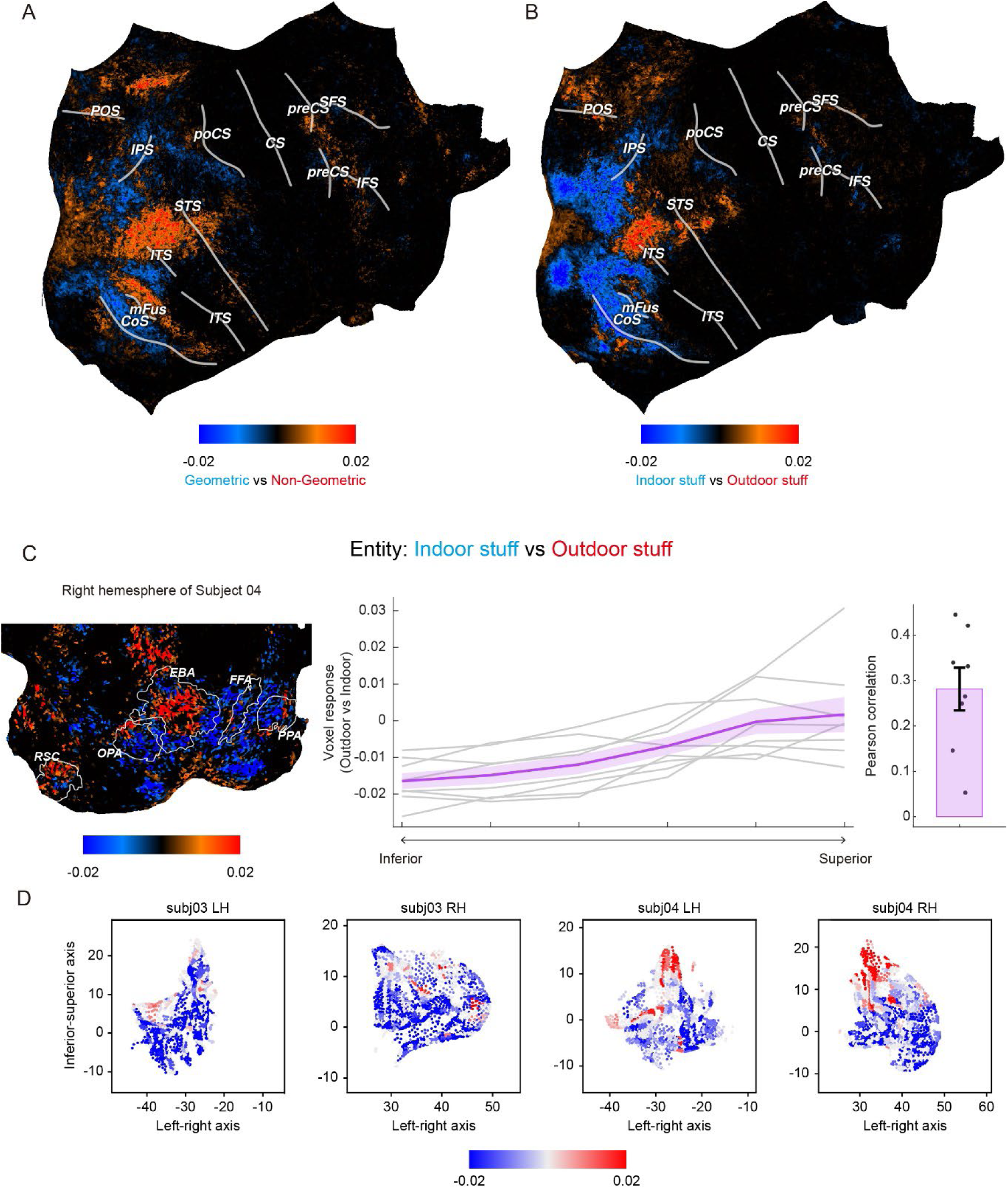
Relation-based and entity-based contrast analyses. (A) Contrast between geometric and non-geometric relation representations. (B) Contrast between indoor-stuff and outdoor-stuff entity representations. Contrast values in (A) and (B) were computed and displayed only for voxels showing significant prediction by the encoding model. (C) Gradient analysis of indoor-stuff and outdoor-stuff representations within the OPA. The left map illustrates the contrast in one representative subject. The middle panel shows contrast values binned along the inferior-superior anatomical axis. The right panel shows Pearson correlations between anatomical position and contrast value for individual subjects. (D) Flattened cortical maps from representative subjects showing the spatial distribution of the indoor-stuff versus outdoor-stuff contrast within the OPA (See Supplementary Figure 3 for all subjects). Point positions reflect anatomical coordinates and colors indicate contrast values.

We next constructed an entity-based contrast between labels belonging to the indoor-stuff and outdoor-stuff categories. Regions near the collateral sulcus, including PPA, showed greater sensitivity to indoor than outdoor scene content, whereas regions near the parieto-occipital sulcus in RSC showed opposite pattern (Figure 6B). Previous studies have shown that the OPA encodes scene layout (Henriksson et al., 2019; Lescroart & Gallant, 2019; Wu & Li, 2026). We further quantified the indoor-outdoor contrast by testing for a representational gradient within the OPA. At the group level, the outdoor-stuff versus indoor-stuff contrast revealed a significant inferior-to-superior gradient, progressing from greater sensitivity to indoor content in inferior OPA to greater sensitivity to outdoor content in superior OPA (*r* = 0.281, *t* = 5.98, *p* <.001; Figure 6C). This pattern in OPA is consistent with previous findings on the cortical organization of 3D scene layout representations (Lescroart, 2026; Lescroart & Gallant, 2019). These results illustrate how the decomposable, context-independent structure of triplet embeddings can generate anatomically specific hypotheses about the cortical organization of scene semantics.

### Triplet encoding differs between macaque IT and human high-level visual cortex

The preceding analyses established behaviorally prioritized triplet embeddings as an interpretable model of relational scene representations in the human visual cortex. We next asked whether the relative contributions of triplet-based semantic features and image-based visual features differed across primate species. The recently released Triple-N dataset contains dense Neuropixels recordings from macaque inferior temporal (IT) cortex obtained while macaques viewed 1,000 natural images from the NSD Shared1000 set, the same images viewed by all NSD subjects (Y. Li et al., 2026). This shared stimulus set enables direct comparisons across species, despite substantial differences in neural measurement. Of the 1,000 images, 412 were also annotated in PSG dataset; all macaque analyses reported below were restricted to this overlapping image subset.

We first compared the predictive performance of VGG19 and the behaviorally prioritized top-3 triplet model in macaque IT and human high-level visual cortex. Previous work found that macaque IT activity was better predicted by features from image-trained ANNs predict than by features derived from LLMs. Here, we fitted separate encoding models based on VGG19 and top-3 triplet models using multiple-kernel ridge regression with cross-validation. For the human analysis, corresponding models were trained on PSG-annotated NSD images and evaluated on the 412 images shared across the NSD, Triple-N, and PSG datasets. In this and all subsequent cross-species analyses, we included only units or voxels with positive prediction correlations for the triplet encoding model. We then used Deming regression to characterize the linear relationship between VGG19 and triplet-model predictive performance. The fitted slope was 0.686 in macaque IT, substantially lower than 0.903 in human high-level visual cortex (Figure 7A), suggesting that triplet features captured less of the VGG19-predictable response in macaques than in humans.

**Figure 7.**
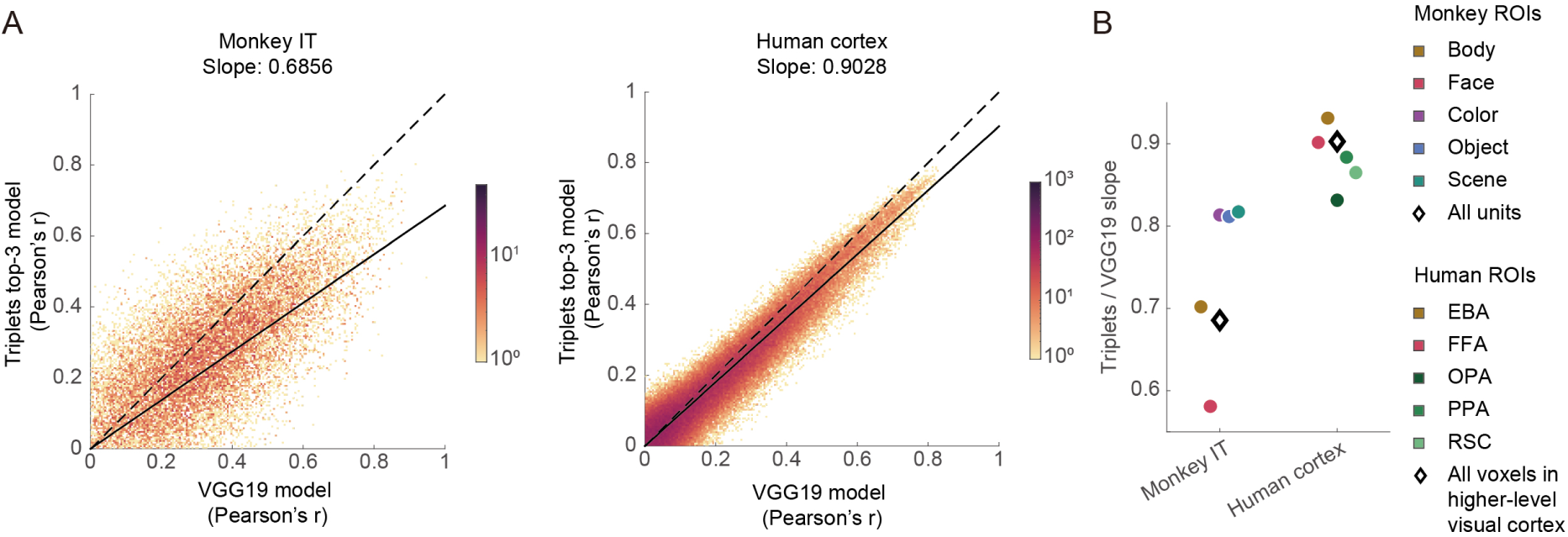
Cross-species differences in the relative encoding of triplet-based features. (A) Scatter density plots showing the relationship between encoding performance for the VGG19 and behaviorally prioritized top-3 triplet models in macaque IT responses from the Triple-N dataset (left) and human high-level visual cortex responses from the NSD dataset (right). Each point represents the log-scaled density of the unit or voxel distribution. Dashed lines indicate equal encoding performance for the two models, and solid lines show the Deming regression fits. (B) ROI-specific Deming regression slopes characterizing the relationship between top-3 triplet and VGG19 encoding performance across macaque and human visual ROIs. Circles represent slopes for individual ROIs. Diamonds represent slopes computed after pooling macaque IT or human high-level visual cortex (including ventral, lateral and parietal streams), respectively.

Analyses within category-and scene-selective regions revealed a further difference in the regional organization of this relative model preference. In macaques, the fitted slopes were lower in body-and face-selective regions than in color-, object-, and scene-selective regions. In humans, by contrast, the slopes in the EBA and FFA were higher than those in the scene-selective ROIs (Figure 7B). This contrasting regional ordering suggests that the cross-species difference did not take the form of a uniform shift across high-level visual cortex. Instead, the difference was particularly evident in regions selective for bodies and faces. One possible interpretation is that macaque IT activity captures visual and local entity co-occurrence statistics but are less strongly aligned with the more complex relational semantics encoded by triplet model. These complex information in triplets may be especially important for body-and face-related representations because interactions involving people and animals often convey actions, intentions, and social relations. The comparatively high slopes in the human EBA and FFA are consistent with this possibility. However, this interpretation remains tentative given differences in recording modality and ROI definition across species.

## Discussion

This study examined how relational information in natural scenes is organized in the human brain. By combining scene-graph representations, behavioral measurements, and large-scale 7T fMRI data, we found that triplet-based representations captured reliable structure in visually evoked cortical activity, and that their correspondence with neural representations improved when greater weight was assigned to relational units that were more perceptually accessible. The decomposable structure of the triplet representation further revealed partially dissociable contributions of entity and relation information, while local triplet content remained aligned with the spatial preferences of high-level visual cortex. Together, these findings suggest that the semantic representation of natural scenes is not a global summary, but retains internal structure shaped by attention, semantic composition, and spatial origin.

### Attentional selection shapes scene triplet representations

Scene triplets provide a structured unit for describing the semantic content of natural images. Unlike isolated object labels, triplets preserve how entities are connected. Unlike free-form verbal descriptions, they remain discrete and local, with each triplet tied to particular entities and image regions. Our behavioral results showed that triplet differed systematically in their accessibility to observers. Triplets involving entities that were larger, more salient, or closer to fixation were judged more accurately, suggesting that access to relational scene information is constrained by visual input and may reflect a bottom-up form of implicit attention. Accessibility was also influenced by relation type and graph properties: geometric relations differed from non-geometric relations, and triplets appearing more frequently were more readily recognized. Geometric relations are common and often specify the stable spatial structure of a scene, whereas non-geometric relations, such as *touching* and *riding*, may convey richer information about action, social interaction, or affordance. Previous work has shown that gaze preferentially samples regions carrying richer scene meaning, consistent the explicit attentional selection in natural scene understanding (Henderson & Hayes, 2017; Murlidaran et al., 2026; Murlidaran & Eckstein, 2025). Our findings suggest a parallel implicit mechanism: observers preferentially process the triplets that carried more interpretable or behaviorally relevant scene meaning, even in the absence of explicit eye movements.

These behavioral regularities provided more than a description of triplet judgments. When used to prioritize triplets in the neural model, they increased correspondence with cortical representations across high-level visual regions. Models incorporating behavioral prioritization predicted visual cortical activity better than the unweighted triplet models, indicating a shared attentional mechanism underlying the perceptual accessibility and the semantic representation in high-level visual cortex. Importantly, the behavioral weighting model was estimated from an independent group of subjects viewing only a subset of the images. Its ability to improve neural predictions across the larger image set therefore suggests that it captured regularities in relational accessibility that generalize across subjects and images.

The cross-species analysis provides an additional heuristic constraint on the interpretation of the model. In humans, the triplet model performed comparably to VGG19 in high-level visual cortex, whereas in macaques it performed worse than VGG19. This pattern suggests that triplet-based semantic information is more closely aligned with human visual cortical responses than with macaque IT responses. It also suggests that the model’s contribution in humans cannot be reduced to generic visual features captured by VGG19. Bodies and faces in natural scenes frequently participate in structured relations involving agency, intention, physical interaction, and social behavior. Such information is explicitly represented in scene triplets but only indirectly available in category-trained image features. In humans, the relational semantics may guide attentional selection toward socially meaningful bodies and faces, increasing the alignment of body-and face-selective regions with triplet representations.

### Cortical organization of decomposable scene semantics

Isolating neural representations of relations in natural scenes is challenging because entities and relations usually occur within the same context. Large language models and artificial neural networks can capture both conceptual and relational information, but these components are difficult to separate because they are often entangled within context-dependent representations. A major advantage of the triplet framework is that the semantic content of a scene can be decomposed into constituent entities and the relations that connect them. Partial RSA revealed partly dissociable cortical distributions: entity-related information was prominent in inferior temporal cortex and regions along the IPS, whereas the LPC contained unique contributions from both entity and relation representations. Voxel-wise encoding models converged on the same pattern, providing complementary evidence that relation-related variance in natural-scene responses cannot be fully accounted for by entity information alone. These results suggest that LPC may represent relations separately from the specific entities they connect, consistent with recent findings using simpler linguistic stimuli (Chen et al., 2026). At the same time, entity-dominant representations observed in the ventral stream align with previous findings on object and concept representation (Grill-Spector & Weiner, 2014; Huth et al., 2016; Popham et al., 2021).

The decomposable feature space also allowed us to move from broad entity-relation separation to more targeted hypotheses about the cortical organization of particular semantic dimensions. Removing entity information and contrasting geometric with non-geometric relations revealed differential sensitivity near lateral parietal and fusiform cortex, regions that also showed substantial gains from behavioral prioritization. Conversely, entity-based contrasts recovered anatomically meaningful distinctions between indoor and outdoor scene content, including an inferior-to-superior gradient within OPA. Although this result should be treated as heuristic, it nevertheless demonstrates how the triplet model can move from general scene selectivity to more specific semantic dimensions in visual cortex. Together, these findings illustrate an important distinction between structured triplet representations and global scene embeddings: once scene meaning is represented in separable components, specific semantic dimensions can be manipulated and linked to anatomically localized neural responses.

Importantly, this decomposability should not be taken to imply that entities and relations are fully independent, either in the RotatE model or in visual cortical representations. In the triplet framework, the two components remain structurally bound. During training, relation embeddings are learned jointly with the embeddings of the entities they connect because the loss function is defined over complete head-relation-tail triplets. This binding is preserved in the behaviorally prioritized model, in which the top-k selection operates on complete triplets rather than on isolated entity or relation labels. Moreover, the overlapping entity-and relation-related effects in the LPC indicate that analytical decomposability does not necessarily correspond to anatomically separate neural systems. The framework therefore permits the contributions of entity and relation to be distinguished analytically while retaining their correspondence within structured relational units.

### Relational scene semantics remain spatially grounded

Retinotopic organization provides a fundamental scaffold for visual processing (Groen et al., 2022). Although triplets encode high-level semantic structure, each triplet remains tied to the image regions occupied by its constituent entities. We found that voxels preferring the left or right visual field were better predicted by triplet models derived from the corresponding side of the image than by models derived from the opposite side. This pattern of result provides direct evidence that semantic information follows retinotopic preferences. Thus, at least along the horizontal visual-field axis, high-level cortex does not represent scene meaning in a completely location-invariant manner but preserves information about where local relational content originates.

The vertical dimension did not exhibit the same pattern of result. Rather than showing an interaction between voxel preference and triplet-model visual field, triplet models derived from the bottom visual field predicted cortical representations more strongly than those derived from the top visual field, regardless of voxel preference. One possibility is that top and bottom visual field preferences are less clearly differentiated in the high-level regions examined here (Groen et al., 2017; Silson et al., 2015). Alternatively, this asymmetry may reflect systematic differences in the amounts or types of semantic information appearing at different vertical positions (Adams et al., 2016; Greene, 2013). The lower portions of natural scenes often contain reachable objects, ground surfaces, animals, bodies, and actions and may therefore contain more of the relational information captured by the triplet model. This interpretation remains speculative. Nevertheless, the contrast between the horizontal and vertical results identifies an important boundary condition: the spatial grounding of relational scene representations may differ across visual-field dimensions.

More broadly, the spatially localized triplet embeddings provide a flexible framework for future investigations of natural scene understanding. Combining this framework with eye tracking could test whether gaze preferentially samples behaviorally prioritized triplets, whereas experimental manipulations of attention could determine how current goals alter the selection of relational information. Memory paradigms could further reveal which entities and relations are retained and how their spatial organization changes during encoding and retrieval. The same approach could also be extended to developmental and clinical populations or to dynamic social scenes, in which relations between entities may be particularly consequential. By representing scene semantics in an explicit, structured, and spatially localized form, the triplet framework offers a means of investigating not only whether the brain represents scene meaning, but also how that meaning is selected, organized, and used.

## Methods

### RotatE model training

#### Natural Scenes Dataset

Natural scene images and fMRI data were obtained from the Natural Scenes Dataset (Allen et al., 2022). NSD contains whole-brain 7T fMRI responses from eight subjects (six females; age range, 19-32 years) who viewed colored natural scene images over 30-40 scan sessions. Each subject viewed 9,000-10,000 images, including a shared set of 1,000 images presented to all subjects. The images were drawn from the Microsoft Common Objects in Context (COCO) database (Lin et al., 2014). Each image was presented for 3 s with a 1 s inter-trial interval, and was scheduled to appear three times per subject. Subjects performed a recognition task in which they indicated whether each image was new or had been presented previously. Images subtended 8.4° x 8.4° of visual angle, with a central fixation dot superimposed on each image.

#### Panoptic Scene Graph dataset

Relational descriptions of the images were obtained from the Panoptic Scene Graph (PSG) dataset (Yang et al., 2022). PSG provides scene-graph annotations based on COCO panoptic segmentations, such that scene-graph nodes correspond to pixel-level panoptic regions rather than only to bounding boxes. The entity vocabulary comprises 133 categories, including 80 thing categories and 53 stuff categories, and the relation vocabulary comprises 56 relation categories. The dataset includes 47,636 images and 275,371 head-relation-tail triplets. Among these images, 28,962 overlapped with the NSD image set and were used to link PSG-derived relational structure to visually evoked neural responses.

#### RotatE vector training

RotatE represents entities and relations in a complex-valued vector space (Sun et al., 2019). For a head-relation-tail triplet *(h, r, t)*, the head and tail entities are represented by complex vectors *e_h_* and *e_t_*, respectively, and the relation is represented as a rotation vector *r* whose elements are constrained to have unit modulus. A triplet is considered plausible when applying the relation-specific rotation to the head vector yields a vector close to the tail vector:

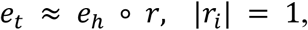

where ∘ denotes element-wise multiplication in complex space. The plausibility of a triplet was quantified by the distance between the rotated head vector and the tail vector:

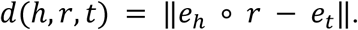

The corresponding RotatE score was defined as the negative distance:

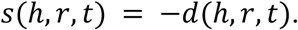

Triplets for which rotated head vectors were closer to the tail vectors therefore received higher scores, whereas less plausible triplets received lower scores.

During training, observed PSG triplets served as positive samples. Negative samples were generated by corrupting positive triplets through replacement of either the head or tail entity. The vectors were optimized with a negative-sampling loss that increased the scores of observed triplets and decreased the scores of corrupted triplets:

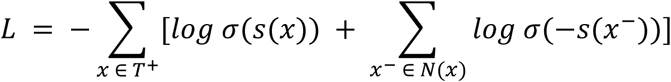

where *T^+^* denotes the set of observed triplets, *N(x)* denotes the negative samples generated for a positive triplet *x*, *s(·)* is the RotatE score, and *σ* is the sigmoid function. This score-based objective encourages the model to learn entity and relation vectors that preserve the relational structure of the PSG annotations.

We trained the RotatE model on the PSG triplets using PyKEEN toolbox. The full set of triplets was divided into training, validation, and test sets containing 80%, 10%, and 10% of the data, respectively. The model used 100-dimensional vectors and was optimized with Adam using a learning rate of 5 x 10^-4^, a batch size of 512, and a maximum of 500 epochs. We applied L2 regularization and used Bernoulli negative sampling during training. Training was monitored on the validation set with early stopping. Optimization was terminated if inverse harmonic mean rank failed to improve by a relative change of 0.002 over 10 consecutive evaluations. The trained model and learned vectors were saved for subsequent image-level triplet representation analyses.

### Behavioral experiment

#### Subjects

Twenty-five healthy adults participated in the behavioral experiment (5 males and 20 females; age range, 18-29 years; M = 21.64, SD = 2.66). All subjects had normal or corrected-to-normal vision and were native Chinese speakers. Subjects provided informed consent and received monetary compensation for completing the experiment. The study was approved by the Committee for Protecting Human and Animal Subjects at the School of Psychological and Cognitive Sciences at Peking University (Institutional Review Board Protocol No. 2025-09-04).

#### Stimuli

The behavioral experiment used 240 natural scene images selected from the subset shared by NSD and PSG. To ensure sufficient scene complexity, images were randomly sampled from PSG-annotated images that contained at least 10 entities and 10 relations. The selected images were then cropped according to the crop coordinates provided by NSD.

Stimuli were presented on a Display++ monitor with a resolution of 1,920 x 1,080 pixels and a refresh rate of 120 Hz. Subjects viewed the images at a viewing distance of 80 cm. Each image subtended 8.4° x 8.4° of visual angle. During image presentation, a red circular fixation marker with a diameter of 10 pixels was superimposed at the center of the image to encourage central fixation.

#### Procedure

Subjects performed a triplet judgment task. On each trial, they first viewed a natural scene image and were then shown three head-relation-tail triplets describing the image. Their task was to identify which semantic component, if any, was inconsistent with the image. Subjects were informed that each triplet contained at most one incorrect component, and that the number of incorrect triplets for a given image could range from zero to three. They were instructed to identify as many incorrect components as possible while minimizing false alarms.

Each trial began with the presentation of a scene image at the center of the screen, with a red fixation marker superimposed at the image center. The image was presented for 3 s, during which subjects were instructed to maintain central fixation. After the image disappeared, three triplets were presented on the screen, with head, relation, and tail components displayed in different colors. Subjects used the mouse to select every semantic component that they judged to be inconsistent with the image. The response window for each image was limited to 8 s.

The experiment consisted of six blocks of 40 images, yielding 240 trials in total. Image order was randomized across the experiment. Before the main task, subjects completed six practice trials to familiarize themselves with the trial structure and response procedure. Eye movements were recorded throughout the experiment using an EyeLink 1000 Plus eye-tracking system at a sampling rate of 1,000 Hz.

Correct triplets were sampled from the PSG annotations for the corresponding image. To construct an incorrect triplet, a triplet other than the correct ones from the current image was used as the base triplet, and exactly one of its three semantic components was replaced. To identify a plausible replacement, we search for a triplet that shared two components with the base triplet but differed in the component to be replaced. Candidate replacement triplets were drawn from other images ordered by their similarities to the current image, beginning with the most similar image and proceeding downward until a suitable candidate was identified. This procedure ensured that each incorrect triplet preserved a plausible a head-relation-tail composition and was also relatively compatible with the overall semantic context of the image, thereby maintained the task difficulty at a reasonable level.

#### Inter-subject consistency analysis

Inter-subject consistency was assessed at the triplet level. For each subject and triplet, the response was coded as an error judgment if the subject selected any of its three semantic components as incorrect. Thus, component-level selections were converted into a binary triplet-level response indicating whether the subject judged the triplet to contain an error.

Response consistency was then calculated for every pair of subjects. For each subject pair, consistency was defined as the proportion of triplets for which the two subjects made the same triplet-level judgment. The inter-subject consistency index was obtained by averaging this proportion across all subject pairs.

Statistical significance was assessed using a permutation procedure. We repeated the pairwise consistency analysis 1,000 times after shuffling subjects’ triplet-level choices to generate a null distribution of consistency values. The observed inter-subject consistency was then compared with this permutation-based null distribution.

#### Triplet feature computation

All features were computed at the level of the whole head-relation-tail triplet. We calculated three classes of triplet-level predictors: visual features derived from the panoptic masks of the constituent entities in each triplet, relational features derived from relation categories and the RotatE vector space, and graph features derived from the scene graph structure of each image.

### Visual features

#### Distance to central fixation

This feature was calculated using the panoptic segmentation masks of the two entities in each triplet. For each entity, we measured the minimum Euclidean distance between the central fixation point and the entity mask. The triplet-level value was defined as the mean of the distances for the two entities.

#### Entity area

This feature was defined as the proportion of the image pixels occupied by the two entity masks in the triplet.

#### Visual saliency

Visual saliency was computed using the Saliency Toolbox (Walther & Koch, 2006). The resulting saliency map was resized to match the image dimensions. Triplet-level saliency was defined as the mean value of all positive saliency pixels within the two entity masks.

For triplets that appeared more than once in the same image, distance to central fixation was summarized by the minimum value across occurrences, whereas entity area and visual saliency were averaged across occurrences.

### Relational features

#### Relation type

Relations were classified as geometric or non-geometric. The geometric category comprised the following 10 relations: *over*, *in front of*, *beside*, *on*, *in*, *attached to*, *hanging from*, *leaning on*, *on back of*, *enclosing*. This classification was based on the semantic meaning of the relation categories and a PCA analysis of the 56 RotatE relation vectors, in which these relations clustered together in the low-dimensional vector space (Figure 2).

#### Triplet score

This feature indexed the semantic plausibility of a triplet. We calculated the score of each triplet using the trained RotatE model and the scoring function described above. Higher scores indicate that the model assigned a higher likelihood to the existence of the triplet.

#### Semantic alignment

This feature quantified the correspondence between an individual triplet and the overall semantic content of the image. We first converted the complex-valued RotatE vectors into real-valued embeddings by concatenating their real and imaginary components. For a triplet *τ = (h, r, t)*, the real-valued head entity embedding, relation embedding, and tail entity embedding were concatenated to obtain a 600-dimensional triplet embedding:

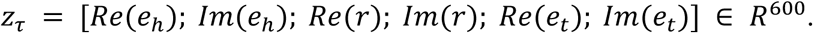

For each image, the embeddings of all annotated triplets were averaged to obtain an image-level embedding:

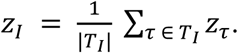

Semantic alignment was then defined as the cosine similarity between the embedding of the current triplet and the image-level embedding:

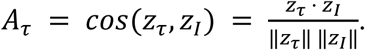

### Graph features

#### Triplet count

This feature was defined as the number of times the same triplet appeared within the image.

#### Triplet degree

This feature quantified the connectivity of the two entities within the scene graph formed by all triplets in the image. The degree of each entity was defined as the total number of incoming and outgoing edges connected to that entity, representing how strongly the entity was connected to the rest of the scene graph. Triplet degree was calculated as the mean degree of the head and tail entities.

#### Logistic mixed-effects regression

To examine the extent to which subjects perceived triplets that were actually present in the scene, the logistic mixed-effects regression was restricted to correct triplets from the image. For each correct triplet, the binary response variable was coded as a false alarm or a correct rejection.

Using a logit link function, we modeled the probability of a false alarm as a function of the triplet-level features described above. Fixed effects included distance to central fixation, entity area, visual saliency, relation type, triplet score, semantic alignment, triplet count, and triplet degree. Relation type was treated as a binary predictor (0 for geometric, 1 for non-geometric), whereas all other predictors were treated as continuous variables. Subject was included as a random intercept to account for between-subject variability in overall response tendency.

### fMRI data analysis

#### MRI data acquisition and preprocessing

Functional MRI data were obtained from the NSD. Data were acquired at 7T using whole-brain gradient-echo echo-planar imaging (EPI), with an isotropic resolution of 1.8 mm and a repetition time (TR) of 1.6 s. Functional images were preprocessed with temporal interpolation to correct for slice timing and spatial interpolation to correct for head motion.

Single-trial response amplitudes were estimated using a general linear model (GLM) approach. We used the NSD third beta version (betas_fithrf_GLMdenoise_RR), estimated with the GLMsingle algorithm (Prince et al., 2022). GLMsingle combines optimized denoising and regularization procedures to improve the estimation of stimulus-evoked voxel responses. Further details are provided in (Allen et al., 2022). Informed written consent was obtained from all participants, and the experimental protocol was approved by the University of Minnesota institutional review board.

#### ROI definitions

We analyzed two sets of regions of interest (ROIs). The first set consisted of category-and scene-selective visual areas, including the fusiform face area (FFA), extrastriate body area (EBA), parahippocampal place area (PPA), occipital place area (OPA), and retrosplenial cortex (RSC). These ROIs were defined based on functional localizers from the NSD experiment.

The second set was derived from the NSD ‘streams’ definitions. The ‘streams’ provides broad subdivisions of visual cortex. These labels are based primarily on fsaverage cortical folding patterns and additionally incorporate NSD noise-ceiling estimates to restrict the ROIs to regions containing reliable stimulus-related signals. We selected the early visual cortex ROI and three higher-level stream ROIs. The early visual cortex ROI corresponds to the union of V1, V2, and V3 subdivisions, whereas the higher-level ROIs correspond to ventral, lateral, and parietal visual streams.

#### Image embedding construction

We constructed image-level embeddings using five models: the unweighted triplet model, behaviorally prioritized top-k triplet model, LLM, object2vec, and ANN. These models provided complementary descriptions of each image for comparison with neural response patterns.

#### Triplet model

The triplet model constructed image embeddings from PSG-annotated triplets using triplet embeddings derived from the trained RotatE model. For each image, we concatenated the real-valued head entity embedding, relation embedding, and tail entity embedding for each triplet, and then averaged the triplet embeddings across all triplets in the image to generate an image-level embedding.

#### Behaviorally prioritized top-k triplet model

The behaviorally prioritized top-k triplet model used the same RotatE triplet embeddings, but retained only the triplets predicted to be most perceptually accessible in each image. Each triplet was assigned a weight equal to its correct-rejection probability predicted by the behavioral logistic regression from distance to central fixation, entity area, visual saliency, and relation type. The weight was computed as:

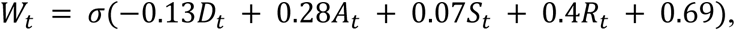

where *D_t_* denotes distance to central fixation, *A_t_* denotes entity area, *S_t_* denotes visual saliency, *R_t_* is the binary code for relation type, and *σ* denotes the sigmoid (logistic) function that maps the linear predictor to a probability between 0 and 1. For each image, triplets were ranked by *W_t_*, and the embeddings of the top-k triplet were combined using a weighted average to obtain the image embedding. We evaluated models with *k* ranging from one to six.

#### LLM

Following Doerig et al. (2025), the LLM-based image embedding was derived from natural-language COCO scene captions. Each NSD image was associated with five human-written captions, each of which was passed through the *all-mpnet-base-v2* sentence-transformer model. The five caption embeddings were averaged to obtain a single image-level embedding. Averaging reduced the influence of caption-specific variability and emphasized semantic information shared across observers.

#### object2vec

Based on Bonner & Epstein (2021), object2vec represents images on the basis of the co-occurrence structure of COCO panoptic object and scene labels. The model extends the logic of word2vec by treating the panoptic segmentation labels within each image as a sentence and learning label vectors using the continuous bag-of-words (CBOW) objective, in which a target label is predicted from the surrounding context labels. We trained object2vec on the 118,287 images in the COCO training set. The CBOW model was trained with a vector dimensionality of 50 for 100 epochs, using negative sampling with 20 negative samples and a window size of 15. After training, each label was represented by a 50-dimensional vector. The image-level object2vec embedding was computed by averaging the vectors of all labels present in the image.

#### ANNs

ANNs transform image pixels through hierarchical stages of visual processing and their representational structures have been suggested to resemble multiple stages of visual processing. We therefore evaluated a set of ANN models and extracted node activations from the pre-readout layer as high-level semantic image representations, except for CLIP models, for which we used the final image embedding rather than the pre-readout layer. Most of these models were also evaluated in Doerig et al. (2025), where their representations were shown to predict neural activity in NSD. We included the following nine ANN models:

- CORnet-S (Kubilius et al., 2018) trained on ImageNet (Deng et al., 2009), obtained from ThingsVision (Muttenthaler & Hebart, 2021).
- AlexNet (Krizhevsky et al., 2012) trained on ImageNet, obtained from Brain-Score (Schrimpf et al., 2018).
- VGG-19 (Simonyan & Zisserman, 2015) trained on ImageNet, obtained from Brain-Score.
- AlexNet-GN trained on ImageNet, obtained from Konkle & Alvarez (2022).
- AlexNet trained with instance-prototype contrastive learning on ImageNet, obtained from Konkle & Alvarez (2022).
- ResNeXt101_32×8d_wsl84 (Mahajan et al., 2018) trained on 914 million public images, obtained from the PyTorch Hub implementation of weakly supervised learning on images.
- NF-ResNet50 trained on ImageNet, obtained from https://github.com/fastai/timmdocs.
- CLIP_RN50_imgs (Radford et al., 2021), corresponding to the visual stream of CLIP with a ResNet-50 backbone, trained on webimagetext.
- CLIP_ViT, corresponding to the visual stream of CLIP with a Vision Transformer backbone, trained on webimagetext and obtained from the OpenAI CLIP repository.

### Representational similarity analysis

We used RSA to quantify the alignment between model representations of natural scene semantics and neural activity patterns in visual cortex. Three complementary analyses were performed: ROI-wise RSA, searchlight RSA, and partial RSA assessing the unique contribution of individual models. For all analyses, voxel-wise beta estimates were averaged across repeated presentations of the same image within each subject, yielding one response estimate per voxel and image. Analyses for each subject were restricted to images shared by NSD and PSG datasets.

#### ROI-based RSA

For each subject and ROI, neural RDMs were constructed in native anatomical space. The dissimilarity between each pair of images was defined as the Pearson correlation distance between their multivoxel response patterns across all voxels within the ROI. Model RDMs were constructed from the corresponding image embeddings using cosine distance. Neural-model correspondence was quantified by the Pearson correlation between the vectorized upper triangles of the neural RDM and the model RDM.

To assess whether each model significantly predicted neural representational geometry, two-tailed t-tests were performed across subjects for each neural-model correlation. False discovery rate (FDR) correction was applied for multiple comparisons across all neural-model comparisons. Model differences within each ROI were tested using one-way ANOVA followed by predefined contrasts.

#### Searchlight RSA

Searchlight RSA was performed in each subject’s native space across all cortical voxels. We moved a spherical searchlight with a radius of 5 mm across the cortical mask. At each searchlight location, activity patterns were extracted for all included images, and a local neural RDM was constructed using Pearson correlation distance between pairs of images. Model RDMs were then compared with the local neural RDM using Pearson correlation, yielding model-alignment values at the center voxel.

Searchlight results were first transformed to surface space and then aligned to fsaverage space for group-level statistical analysis. Group-level significance was assessed using two-tailed t-tests on each fsaverage vertex across NSD subjects. Multiple comparisons were corrected using the Storey procedure for controlling the false discovery rate (Storey, 2002), with alpha = 0.05.

#### Partial RSA for unique model contributions

Partial RSA was used to estimate the unique contribution of each model after controlling for the other model representations. In this analysis, the target model RDM was partially correlated with the neural RDM while the RDMs of the remaining models were included as covariates.

We applied this procedure in two contexts. First, to test whether the top-3 triplet model captured information not explained by other semantic models, we included four model RDMs: top-3 triplet model, VGG-19, MPNet, and object2vec. For each target model, its partial correlation with the neural RDM was computed while controlling for the other three model RDMs.

Second, we used partial RSA to examine the separate contributions of entity and relation information within the top-3 triplet model. For each triplet, the head and tail entity embeddings were concatenated to obtain a 400-dimensional entity embedding. Image-level embeddings for entity and relation were then constructed separately using the same top-3 weighting procedure described above. We constructed separate entity and relation RDMs from these image-level embeddings and assessed their partial Pearson correlations with the neural RDM, allowing us to estimate the unique contribution of entity versus relation information.

### Encoding model

We performed the encoding model analyses using the Himalaya toolbox (Dupré La Tour et al., 2022). We trained a linear encoding model for all cortical voxels in each subject’s native space using banded ridge regression. The model predicted each voxel’s response to the images from the image-level embeddings, formalized as *y = Xw*, where *y* denotes the voxel response vector across images, *X* denotes the image-by-feature design matrix, and *w* denotes the learned feature weights. To reduce the number of feature channels, the head, relation, and tail components of the image-level embeddings for all PSG images were separately reduced using PCA. For each component type, we retained the minimum number of principal components required to explain 95% of the variance, resulting in 48 head components, 18 relation components, and 56 tail components.

Banded ridge regression estimates separate regularization hyperparameters for different feature spaces (Dupré La Tour et al., 2025). We grouped the head and tail components into an entity feature space and treated the relation components as a separate features space, as entities and relations play distinct roles in the RotatE representation.

For each subject, 90% of the images were randomly assigned to the training set and the remaining 10% were held out as an independent test set. Within the training set, five-fold cross-validation was used to select the optimal regularization hyperparameters separately for each feature space and voxel. Hyperparameter candidates were generated through random search. Specifically, 1,000 normalized hyperparameter candidates were randomly sampled from a Dirichlet distribution and then scaled by 10 logarithmically spaced values ranging from 10^-5^ to 10^5^. For each voxel, the selected hyperparameters were those that minimized the squared error (L2) loss between the predicted and recorded responses during the five-fold cross-validation. The selected hyperparameters for each subject and voxel were then used to fit the encoding model on the full training set. Model performance was evaluated on the held-out test set by computing the Pearson correlation between the predicted and observed voxel responses across images for each voxel.

Statistical significance was assessed at both the individual-subject and group levels. At the individual-subject level, voxel-wise significance was evaluated using 1,000 permutation iterations. In each iteration, neural responses in the test set were shuffled across stimuli, and the correlation between predicted responses and shuffled responses was recalculated. The resulting distribution provided a voxel-specific null distribution for the observed prediction accuracy. At the group level, voxel-wise model performance was computed in each subject’s native space, transformed to surface space, and aligned to fsaverage space. Group-level significance was assessed using two-tailed t-tests across NSD subjects, with multiple comparisons corrected using the Storey procedure for controlling the false discovery rate at alpha = 0.05.

To decompose the prediction performance of the full encoding model into entity and relation contributions, we calculated the split-prediction performance generated by each feature space. Specifically, we first obtained the split prediction produced by feature space *i*, and then computed its normalized dot product with the observed voxel response:

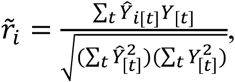

where *Y*_[*t*]_ is the observed voxel response to test image *t*, *Ŷ*_[*t*]_ is the prediction of the full model, and *Ŷ _i_*_[*t*]_ is the split prediction contributed by feature space *i*. This approach has been used in previous encoding-model studies (Chen et al., 2024; St-Yves & Naselaris, 2018) and discussed in work on variance and feature-importance decomposition (Dupré La Tour et al., 2022; Grömping, 2015).

### Encoding-model-based contrast analysis

The encoding model allowed us to predict the responses of individual voxels to arbitrary triplets, as well as to isolated entity or relation components. We therefore used the fitted encoding model to conduct two controlled *in silico* contrasts between selected entity or relation categories.

#### Geometric versus non-geometric relations

The geometric category included the 10 relations defined in the behavioral analysis: *over*, *in front of*, *beside*, *on*, *in*, *attached to*, *hanging from*, *leaning on*, *on back of*, *enclosing*. The non-geometric category included the same number of relations as the geometric category and was selected because these relations pointed in the opposite direction from the geometric relations on the first or second PCA dimension, including *standing on*, *sitting on*, *lying on*, *looking at*, *walking on*, *holding*, *carrying*, *swinging*, *pulling*, and *catching*. To compute *in silico* neural responses for each relation, we projected the relation embedding into the PCA-reduced relation space, concatenated it with zero vectors in the head and tail entity spaces, and applied the fitted encoding weights to obtain a predicted voxel response.

#### Outdoor versus indoor entities

The outdoor category included *mountain*, *river*, *grass*, *gravel*, *road*, *pavement*, *platform*, *fence*, whereas the indoor category included *wood wall*, *tile wall*, *door*, *rug*, *cabinet*, *curtain*, *shelf*, *mirror*. For each entity, we projected its embedding into the PCA-reduced head and tail spaces. The head-space embedding was multiplied by the proportion with which the entity appeared as a head in the PSG dataset. The tail-space embedding was multiplied by the proportion with which the entity appeared as a tail. The two weighted embeddings were concatenated with a zero vector in the relation space. The resulting feature vector was passed through the fitted encoding weights to obtain a predicted voxel response.

### Cortical representational gradient analysis

Cortical representational gradients were analyzed at the vertex level on each subject’s native surface space. Anatomical position of each vertex on the cortical surface was defined using the three-dimensional coordinates of the ‘fiducial’ surface generated from FreeSurfer reconstruction. The ‘fiducial’ surface is defined as the midpoint between the white-matter surface and the pial surface, and therefore preserves the three-dimensional anatomical structure and spatial position of the cortex.

With OPA, we quantified the outdoor-versus-indoor cortical gradient by computing Pearson correlations between each vertex’s anatomical coordinates and its outdoor-versus-indoor contrast value.

We included vertices for which the entity component of the encoding model significantly predicted vertex activity (p < 0.05, uncorrected). Multiple-comparison correction was not applied to this vertex-selection step because the contrast analysis used split predictions from only the entity component of the model, which reduced overall model performance. We therefore used a more liberal threshold to retain a sufficient number of vertices for estimating the outdoor-versus-indoor cortical gradient.

### Comparative analysis of NSD fMRI and the Triple-N dataset

#### Triple-N dataset

The Triple-N dataset (Y. Li et al., 2026) extends the NSD framework to nonhuman primates and contains dense electrophysiological recordings acquired while five adult macaques passively viewed the 1,000 images in the NSD shared1000 set. Neural activity was recorded using Neuropixels NHP probes from early visual areas and inferotemporal cortex, including fMRI-localized category-selective regions. The present analyses focused on the IT recordings, comprising 21,214 visually responsive units distributed across 10 IT ROIs. We used the preprocessed data released by the authors. For each unit, neural response measures were the provided mean firing rate for each image presentation. The study was approved by the Peking University Animal Care and Use Committee (id: Psych-BaoPL-1).

#### Macaque encoding analysis

The encoding analyses were restricted to the 412 images shared by the NSD shared1000 set and the PSG dataset. We compared two feature models: the behaviorally prioritized top-3 triplet model and the pre-readout activations of VGG19. The same random split was used for both models, with 90% of the images assigned to the training set and the remaining 10% held out as the test set.

For each IT unit, we used kernel ridge regression to predict its time-averaged firing rate from the VGG19 features. In its dual form, kernel ridge regression operates on a sample-by-sample kernel matrix rather than directly estimating a coefficient for every feature, making regularized estimation computationally efficient when the feature dimensionality exceeds the number of samples. The regularization parameter α was selected separately for each unit using five-fold cross-validation on the training set from 41 logarithmically spaced values ranging from 10^-5^ to 10^5^.

For the top-3 triplet model, we fitted a banded kernel ridge regression model directly to the triplet image-level embeddings. Because the number of available images was limited, no PCA reduction was applied. Separate regularization hyperparameters were estimated for the entity feature space, comprising the head and tail components, and the relation feature space. Hyperparameters were selected by five-fold cross-validation on the same training set. Specifically, 1,000 normalized feature-space weight vectors were sampled from a Dirichlet distribution and scaled by 41 logarithmically spaced regularization values ranging from 10⁻⁵ to 10⁵. The selected hyperparameters were then used to refit each voxel’s model on the full training set.

Encoding performance was evaluated on the held-out test set by computing, for each IT unit, the Pearson correlation between the predicted and observed time-averaged firing rates across images. To quantify the relative bias toward VGG19 versus top-3 triplet model, we fitted the linear relationship *y = βx* using Deming regression, where *x* and *y* denote the test-set prediction correlations of the VGG19 and top-3 triplet encoding models, respectively. Only units with a positive prediction correlation for the top-3 triplet model were included in this comparison.

#### Human encoding analysis

Human encoding models were fitted separately in each subject’s native space and restricted to voxels in the ventral, lateral, and parietal streams. The 412 PSG-annotated images in the NSD Shared1000 set served as the test set. All remaining PSG-annotated NSD images available for each subject were used for training. VGG19 pre-readout features were fitted using ridge regression, with α selected for each voxel by five-fold cross-validation from 10 logarithmically spaced values between 10⁻⁵ and 10⁵. The top-3 triplet model used the original, non-PCA embeddings and the same banded ridge procedure described above, with separate regularization for entity and relation features.

Prediction performance was quantified by the Pearson correlation between predicted and observed responses across test images. Voxels with positive prediction correlations for the top-3 triplet model were pooled across subjects. The relationship between VGG19 and top-3 triplet performance were estimated using Deming regression.

#### ROI analysis

For each macaque and human ROI, all eligible units or voxels were pooled and the top-3 triplet-to-VGG19 slope was estimated using the same Deming regression described above. Macaque recording ROIs were grouped by functional category into five regions: body (anterior body, AB, and middle body, MB), face (anterior face, AF and middle face, MF), scene (lateral place patch, LPP, and posterior inferotemporal place patch, PITP), color (anterior medial color area, AMC, and central lateral color area, CLC), and object (anterior object, AO and middle object, MO). Human ROI analyses were conducted separately for FFA, EBA, PPA, OPA, and RSC.

## Acknowledgments

This study was supported by grants from STI2030-Major Projects (2021ZD0200204) and the National Natural Science Foundation of China (32271104).

## Supplementary Tables

**Supplementary Table 1.** Logistic mixed-effects model predicting false alarms from visual, relational, and graph-based triplet features. SE: standard error.

| Name | $\beta$ | SE | t | p |
| --- | --- | --- | --- | --- |
| Intercept | -0.7091 | 0.0692 | -10.2406 | 0.0000 |
| Relation type (0 for geometric) | -0.3480 | 0.0577 | -6.0343 | 0.0000 |
| Distance from fixation | 0.1239 | 0.0221 | 5.5974 | 0.0000 |
| Entity area | -0.2922 | 0.0260 | -11.2469 | 0.0000 |
| Visual saliency | -0.0699 | 0.0206 | -3.3966 | 0.0007 |
| Triplet count | -0.1277 | 0.0284 | -4.4929 | 0.0000 |
| Triplet degree | 0.0510 | 0.0264 | 1.9313 | 0.0535 |
| Triplet score | -0.0266 | 0.0257 | -1.0374 | 0.2996 |
| Semantic alignment | 0.0091 | 0.0249 | 0.3657 | 0.7146 |

**Supplementary Table 2.** Reduced logistic mixed-effects model predicting false alarms from visual features and relation type.

| Name | $\beta$ | SE | t | p |
| --- | --- | --- | --- | --- |
| Intercept | -0.6907 | 0.0683 | -10.1080 | 0.0000 |
| Relation type (0 for geometric) | -0.4026 | 0.0528 | -7.6307 | 0.0000 |
| Distance from fixation | 0.1339 | 0.0218 | 6.1485 | 0.0000 |
| Entity area | -0.2843 | 0.0249 | -11.4016 | 0.0000 |
| Visual saliency | -0.0734 | 0.0205 | -3.5786 | 0.0003 |

## Supplementary Figures

**Supplementary Figure 1.**
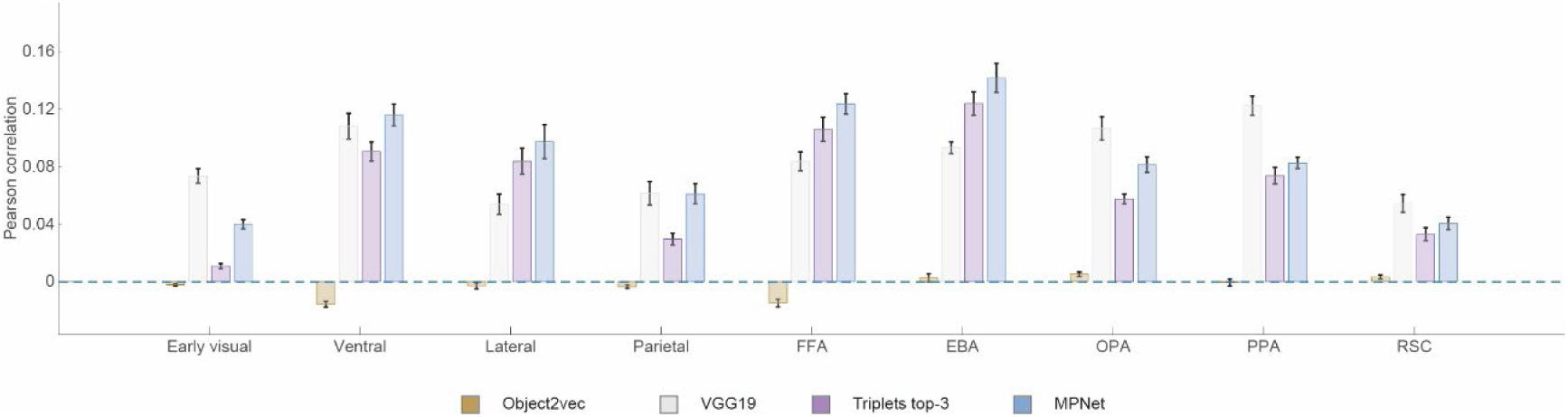
The partial correlations between each model RDM (Object2Vec, VGG19, top-3 triplet model, and MPNet) and the neural RDMs after controlling for the other three model RDMs. Error bars indicate the SEM across participants.

**Supplementary Figure 2.**
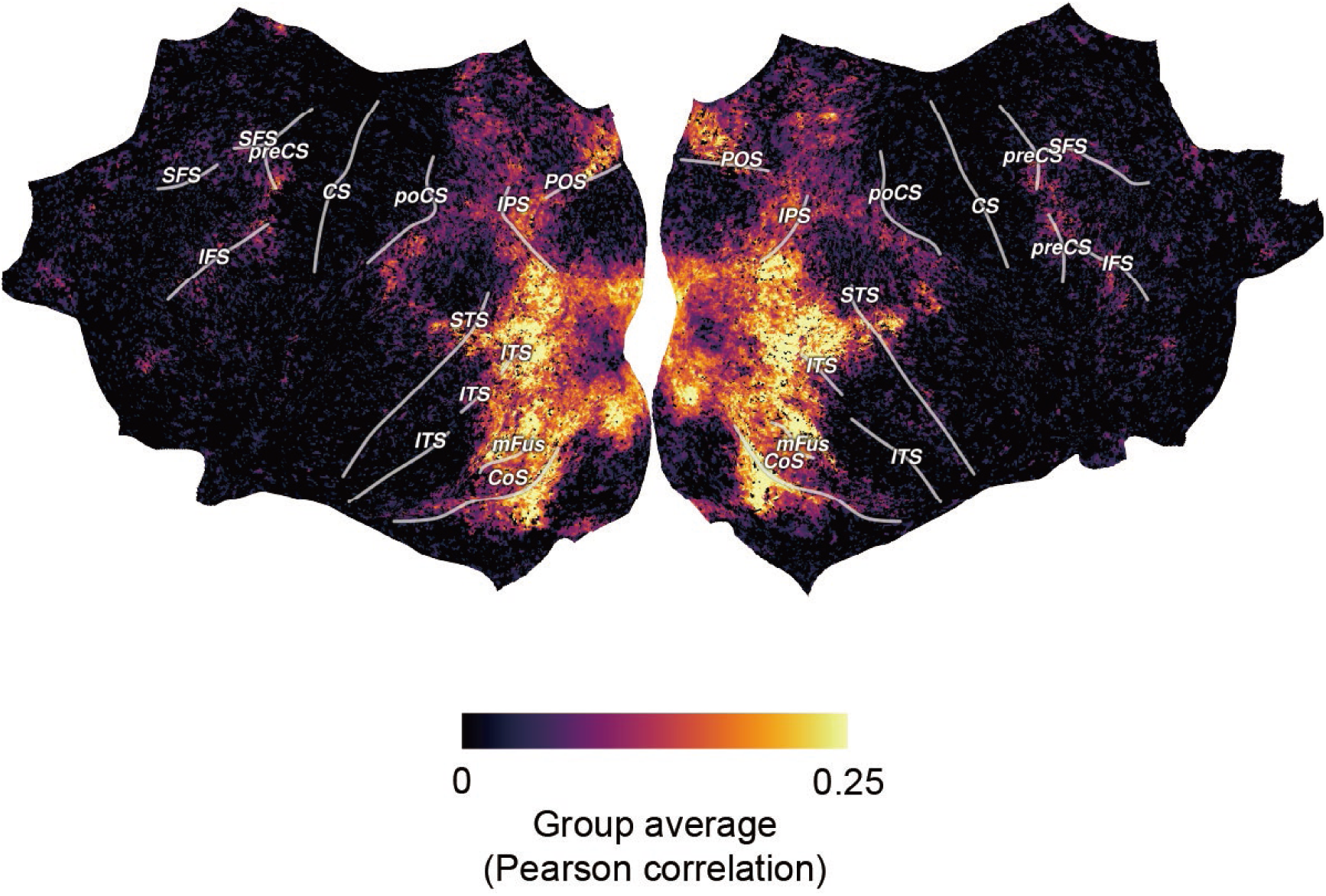
Encoding model performance. Cortical maps show the group-averaged Pearson correlations between model-predicted and observed neural responses in the held-out test set. Statistical significance was assessed using one-tailed *t*-tests against zero with Storey’s FDR correction. Nonsignificant vertices are shown in black.

**Supplementary Figure 3.**
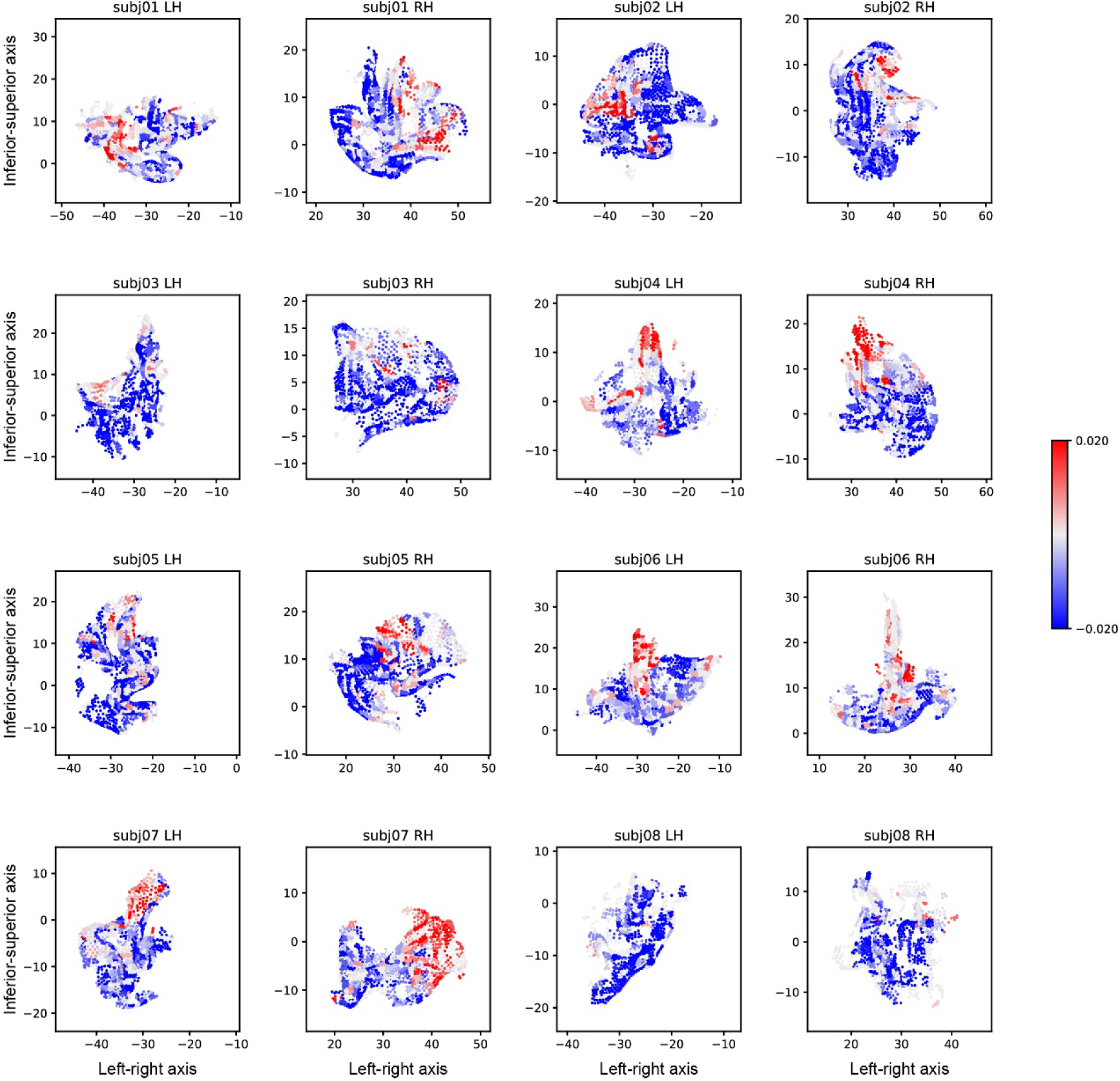
Flattened cortical maps from all participants showing the spatial distribution of the indoor-stuff versus outdoor-stuff contrast within the OPA. Red vertices indicate a preference for outdoor stuff, whereas blue vertices indicate a preference for indoor stuff.

